# Direct Measurement of Unsteady Microscale Stokes Flow Using Optically Driven Microspheres

**DOI:** 10.1101/2020.10.28.354738

**Authors:** Nicolas Bruot, Pietro Cicuta, Hermes Bloomfield-Gadêlha, Raymond E. Goldstein, Jurij Kotar, Eric Lauga, François Nadal

## Abstract

A growing body of work on the dynamics of eukaryotic flagella has noted that their oscillation frequencies are sufficiently high that the viscous penetration depth of unsteady Stokes flow is comparable to the scales over which flagella synchronize. Incorporating these effects into theories of synchronization requires an understanding of the global unsteady flows around oscillating bodies. Yet, there has been no precise experimental test on the microscale of the most basic aspects of such unsteady Stokes flow: the orbits of passive tracers and the position-dependent phase lag between the oscillating response of the fluid at a distant point and that of the driving particle. Here, we report the first such direct Lagrangian measurement of this unsteady flow. The method uses an array of 30 submicron tracer particles positioned by a time-shared optical trap at a range of distances and angular positions with respect to a larger, central particle, which is then driven by an oscillating optical trap at frequencies up to 400 Hz. In this microscale regime, the tracer dynamics is considerably simplified by the smallness of both inertial effects on particle motion and finite-frequency corrections to the Stokes drag law. The tracers are found to display elliptical Lissajous figures whose orientation and geometry are in agreement with a low-frequency expansion of the underlying dynamics, and the experimental phase shift between motion parallel and orthogonal to the oscillation axis exhibits a predicted scaling form in distance and angle. Possible implications of these results for synchronization dynamics are discussed.

## I. INTRODUCTION

In his landmark 1851 paper on viscous fluids [1], George Gabriel Stokes not only developed the theoretical framework for understanding the competition between inertial and viscous forces, but he also considered several physical situations in which that competition is particularly simple to analyze. These include his celebrated problems I and II — viscous fluid in the half space adjacent to a no-slip wall that is impulsively started into motion or oscillated from side to side at some frequency *ω* — as well as the case of a sphere oscillated back and forth. From these oscillatory cases and his newly identified ‘index of friction’ (what we now term the kinematic viscosity *v* = *η/ρ_f_*, *η* and *ρ_f_* being the dynamic viscosity and density of the fluid), he identified from the diffusion equation *u_t_* = *_v_xx__*, for a component *u* of the fluid velocity, the viscous penetration length

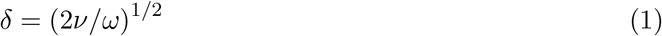

as the distance over which oscillatory motions decay away from the driving surface. Furthermore, the fluid oscillations at some distance *r* from the driving body are phase shifted relative to the drive by an angle proportional to *r/δ*.

There is, of course, no doubt of the *validity* of his analysis of these particular problems. Yet, in the motivating biophysical context we consider here there has been longstanding uncertainty about the *relevance* of unsteadiness to phenomena that are strongly in the Stokes regime, such as the beating of eukaryotic flagella and the motion of tracer particles in flows driven by flagellated organisms. For example, in models for ciliate propulsion [2], it has been recognized that around large microorganisms covered in a dense cilia carpet, unsteady effects are significant within a region near the organism surface of width ~ *δ,* and outside the flow may be considered steady. Consider for example the well-studied multicellular organism alga *Volvox* [3], a spheroid of radius ~ 200 *μ*m, covered with thousands of biflagellated somatic cells each 10 *μ*m in diameter, spaced some 20 *μ*m apart, whose flagella of length *ℓ* ~ 25 *μ*m beat at a frequency *f* ~ 25 Hz. The viscous penetration depth *δ* ~ 110 *μ*m is significantly less than the organism’s circumference, but it is intriguingly close to the wavelength of metachronal waves that *Volvox* exhibits [4, 5]. These are long-wavelength phase modulations of the beating in the form like that of a stadium wave which, in *Volvox,* have a wavelength ~ 100 *μ*m [5]. Thus, even nearest-neighbor somatic cells have a significant phase shift. This is in contrast to the situation in *Chlamydomonas,* the unicellular relative of *Volvox* whose size is comparable to *Volvox* somatic cell and whose two 10 — 12 *μ*m flagella are mounted just a few microns apart and beat at ~ 50 Hz; the phase shift between the flagella is indeed rather small. Yet, nearly all models of flagellar synchronization and in particular of metachronal wave formation [6–14] assume as a starting point the *steady* Stokes equation. It is only recently, in the context of the dynamics of tracer particles in flows generated by the beating flagella of alga [15–18] that unsteadiness has been identified as a potentially significant feature of biophysical flows.

With the goal of motivating further studies of these phenomena, we introduce an experimental setup by which optical trapping methods [19, 20] are used to measure the motion induced by unsteadiness over a broad angular sector around a central oscillated microsphere. This setup allows for a precise test of the underlying microhydrodynamic theory, with results that are complementary to recent experimental studies of oscillatory flows driven by the more complex beating of *Chlamydomonas* flagella, where the phase lag experienced by tracer particles was measured directly [16, 18]. As shown in earlier work on synchronization [21], the far-field flows due to eukaryotic flagella are accurately represented by moving point forces. Thus, we expect the present results to inform future analysis of flagellar interactions on the basis of simplified representations of their dynamics.

We begin in Sec. II with a description of the experimental setup, the frequency response of the optical trap used to oscillate a microsphere surrounded by an array of tracers, and a discussion of inertial corrections to the motion of the microspheres. The results presented in Sec. III comprise the motion of tracer particles at varying distances and angular positions relative to the driven microsphere. The theoretical analysis of their Lagrangian dynamics is done with the Eulerian flow field of the classical solution for motion around an oscillating sphere. A low-frequency expansion, valid when δ is large compared to distances from the central sphere, is used to obtain a geometrically simple result for the tracer trajectories, which are elliptical Lissajous figures. As the tracers are submicron, they exhibit substantial thermal fluctuations which compete with the deterministic displacements from the oscillating flow, and this can be quantified by a suitable Péclet number that varies with oscillation frequency and distance from the central particle. The phase shift between motion along the two Cartesian directions, which is responsible for the shape of those orbits, is calculated in the low-frequency limit and found to be in excellent agreement with the data. The implications of the observed phase shift between the driven and tracer particles on synchronization processes relevant to the biomechanics of cilia are discussed in the concluding Section IV.

## II. SETUP AND EXPERIMENTAL METHODS

A large silica microsphere (radius *a*_0_ ~ 2.77 *μ*m, *ρ*_0_ = 2.65 × 10^3^ kg m^−3^) is forced to oscillate horizontally along the *x*-axis with an amplitude *ξ*_0_ = ξ_0_ exp(i*ωt*) in water (density *ρ_w_* = 10^3^ kg m^−3^, viscosity *η* = 10^−3^ Pas) by means of optical tweezers. Smaller passive polystyrene microspheres (radius *a*_1_ = 0.505 *μ*m, density *ρ*_1_ = 1.05 × 10^3^kgm^−3^), also referred to as *probes* or *tracers,* are located in the horizontal (*x, y*)-plane at three different distances *R_i_* (*R*_1_ = 9.5118 *μ*m, *R*_2_ = 15.853 *μ*m and *R*_3_ = 25.365 *μ*m) from the central sphere, and ten different angles *θ_j_* (*j* = 1 ⋯ 10) equally spaced within the interval [0, *π*], as shown in Fig. 1. The Lagrangian displacement of a polystyrene sphere located at (*R_i_*, *θ_j_*) due to the flow generated by the central bead is denoted by 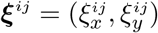.

**FIG. 1.**
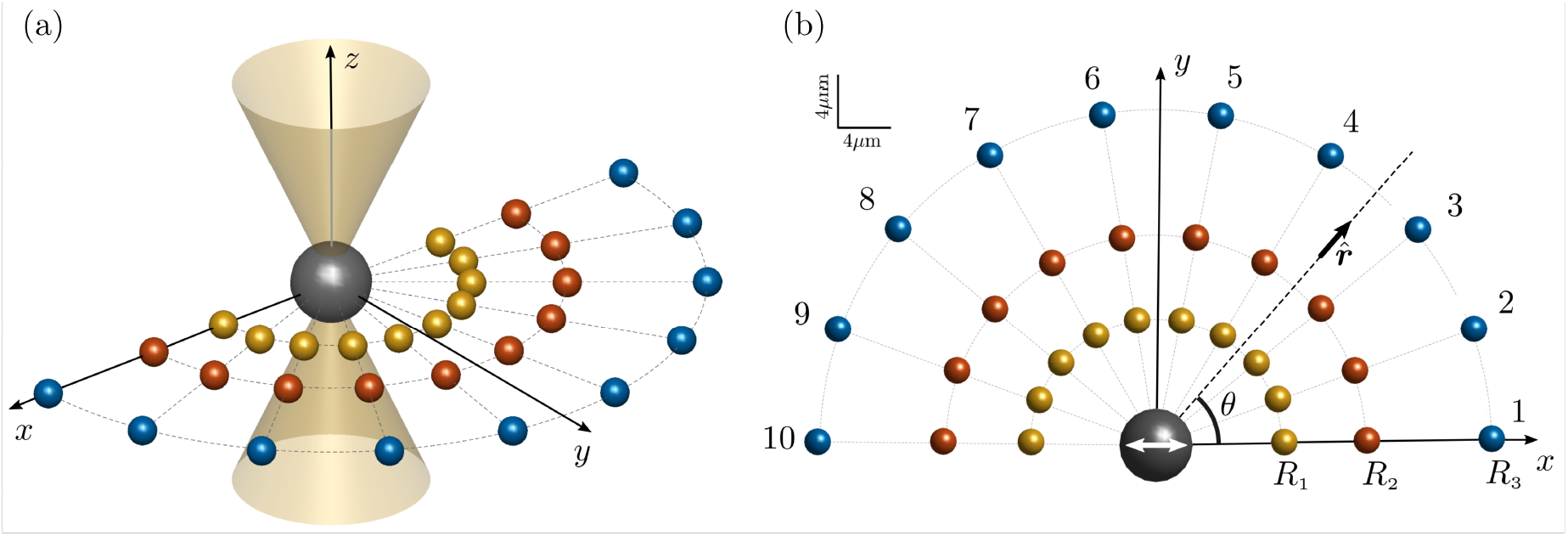
Schematic of the experiment. (a) The large silica microsphere (grey) of radius *a*_0_ = 2.77 *μ*m is oscillated at various frequencies by means of an optical trap. Previous to the actuation of the large particle, passive microspheres (yellow, red, blue) with radius = 0.505 *μ*m are placed at three different distances *R*_1_ = 9.5118 *μ*m, *R*_2_ = 15.853 *μ*m and *R*_3_ = 25.365 *μ*m from the center of the driving bead. They are held in position *via* a multiple trap system, and released automatically upon actuation of the central sphere, whose motion along the *x*-axis is shown by a double white arrow.

The particles used in each experimental run were extracted from dilute suspensions of silica (Bangs Laboratories) and polystyrene (Polyscience) microspheres. The polystyrene and silica beads were sufficiently dilute that no particles other than those used as oscillator or probes interfered with the laser beam during an experiment. The solution was sealed between a microscope slide and a coverslip separated by a 150 *μ*m gap and held together by NOA 68 UV-cured glue. Microspheres were trapped at least 50 *μ*m from the chamber walls to minimize any wall-particle interactions.

The tweezers setup is as described elsewhere [14, 22, 23]. Briefly, the beam of a diode-pumped solid-state laser (CrystaLaser IRCL-2W-1064, 1064 nm wavelength, 2 W maximum output power) is deflected by a pair of acousto-optic deflectors (AA.DTS.XY-250@1064 nm, AA Opto-Electronic) and directed to the back illumination port of a Nikon Ti-E inverted microscope, in which the beam is reflected by a dichroic mirror and focused on the sample by a Nikon Plan Apo VC 60x objective (NA = 1.20). The samples were viewed with brightfield illumination and the dynamical response of the microspheres to the oscillating flow was recorded by high speed camera (Phantom V5.1) at 25,000 frames per second. The acousto-optic deflector allows for time-sharing of the laser beam so that multiple traps located at prescribed positions can be created in the (*x, y*) focal plane. The stiffness of the trap for the silica when it is trapped alone was determined to be *k* = 50 ± 1 pN *μ*m^−1^, by measuring the standard deviation of the particle’s thermal fluctuations.

Initially, the silica bead is trapped at the origin of the coordinate system while the ten polystyrene particles are held at locations (*R_i_,θ_j_*), one *R_i_* at a time. The central particle is driven by moving its optical trap along the *x*-axis by sampling the path as *N_p_* points which are cyclically visited by the acousto-optic deflector. Once the polystyrene particles are released, the entire laser power is reassigned to the oscillating trap and the trajectories of the silica and polystyrene particles are recorded. The corresponding tracks in the (*x, y*)-plane are extracted using a bespoke image segmentation tracking algorithm. The system was optimized to reach driving frequencies up to 400 Hz for a trap oscillation amplitude of 2.15 *μ*m. The oscillation frequency and amplitude are limited by the dynamics of the particle (see below) and the finite size of the optical potential well. Once they are optimized, the maximal *N_p_* for path sampling is obtained from the highest target frequency and a time-sharing frequency of 20 kHz set by the bespoke electronics that control the acousto-optic deflector; *N_p_* = 50 in experiments reported here. Under such conditions, the drive bead follows a sinusoidal pattern, even when it does not remain close to the trap center.

The dynamics of a microsphere forced by an optical trap whose position is laterally oscillated is a well-studied problem [24]. When, as is the case here, the displacement of the driven particle is sufficiently small that the optical force exerted on the sphere is linearly proportional to the distance from the axis of the beam, and the trap and particle positions oscillate as *ζ*_0_e^*iωt*^ and ζ_trap_e^*iωt*^, then momentum conservation in the *x*-direction takes the form

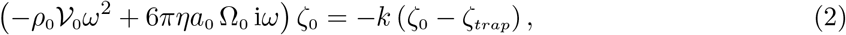

where the left hand side represents the inertia of the particle itself (whose volume is 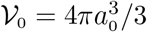) and the drag force, while the right hand side is the trap force. The drag force, found in the original derivation by Stokes in 1851 [1] and in more modern treatments [25, 26] has a factor Ω_0_ that corrects the familiar zero Reynolds number Stokes drag for fluid inertia,

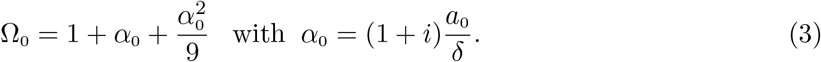

From (2) and (3), we identify three characteristic times scales associated with the experiment, the shortest of which is that for fluid momentum to diffuse on the scale of particle,

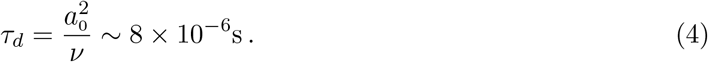

Next is that for inertial oscillations of the sphere in the trap,

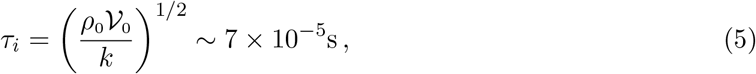

and finally the time scale over which a particle viscously relaxes to the trap center,

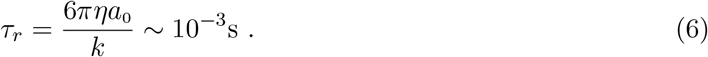

With these definitions, we have

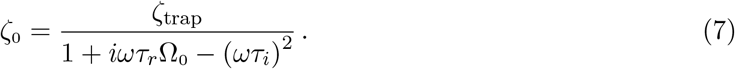

At the highest frequencies probed (400 Hz), *ωτ_i_* ~ 0.18 and particle inertia contributes only a few percent to the response of microspheres. Similarly, from (3) we deduce that the maximum contribution of momentum diffusion has *ωτ_d_* ~ 0.02, so |*α*_0_| ~ 0.14, yielding a modest correction to the force amplitude Ω_0_, and higher-order contributions are negligible. By the quadratic scaling of Td with sphere radius, this simplification is due to the use of microspheres. The complex structure of (7) implies that there is a phase shift between the trap and the driven particle, but as we are interested in the response of the tracers relative to the driven microsphere, we ignore that phase shift and use the motion of the driven bead as the time reference in the following and define ξ_0_(*ω*) = |ζ_0_1; we adopt a time origin such that the driven particle’s position is ξ_0_ cos(*ωt*). Neglecting particle inertia and quadratic terms in Ω_0_, we have

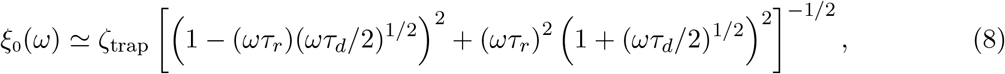

which represents only minor deviations from the Lorentzian form [1 + (*ωτ_r_*)^2^]^−1/2^. This ratio and its Lorentzian approximation are plotted in Fig. 2 a as a function of the rescaled frequency *ωτ_r_*. With no free parameters the agreement between the prediction of (8) and experiment is excellent. Figure 2b shows the accurately sinusoidal displacement of the oscillated particle.

**FIG. 2.**
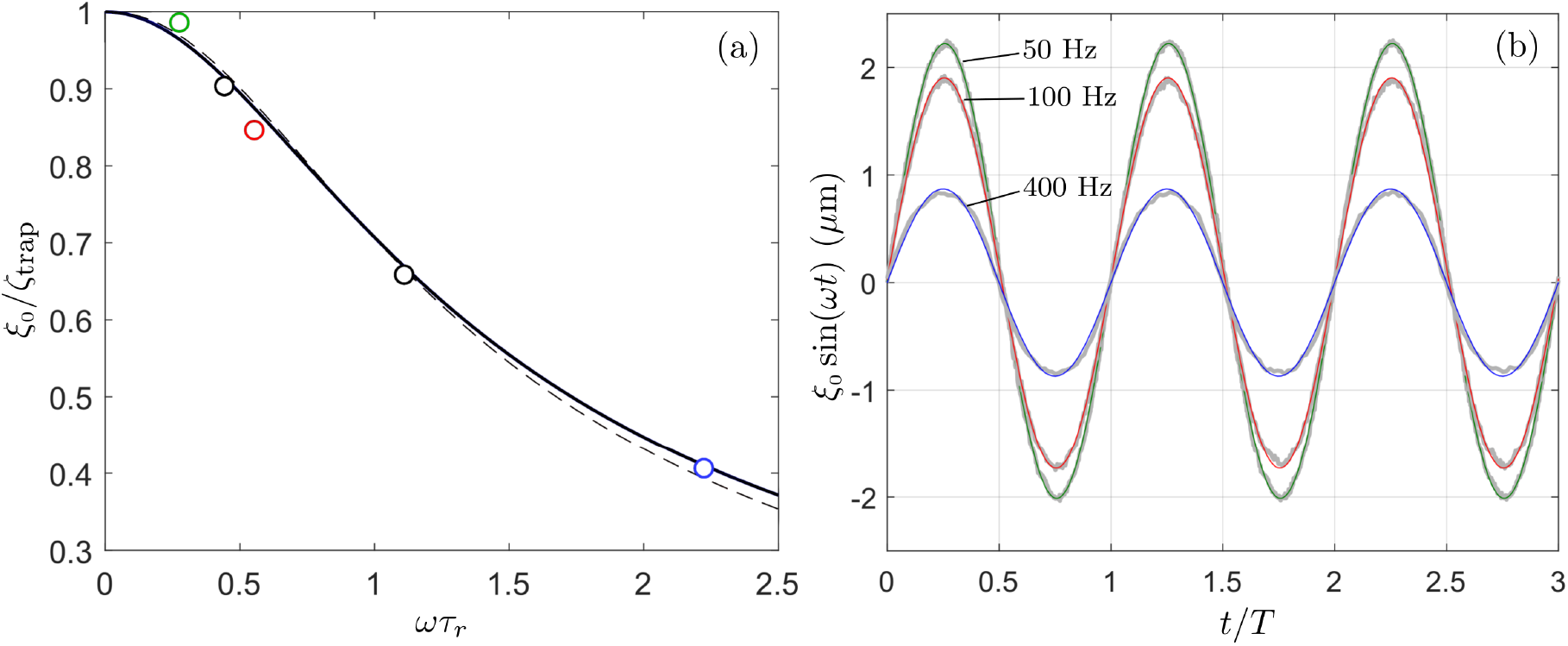
Oscillations of the driven microsphere. (a) Amplitude of oscillations relative to trap oscillation amplitude as a function of frequency. The theoretical prediction in Eq. 8 with *τ_d_* = 0 (solid blue line) matches well the experimental data (open circles). Dashed red line indicates small correction obtain by including finite *τ_d_* corrections. (b) Measured *x*-position of the bead as a function of the time (thick gray lines), fitted with a sinusoidal function (colored lines) at three frequencies. For clarity of presentation, the origin of time has been shifted to align the different curves. The good match between the experimental profiles and the fitting functions validates the linear approach used to derive Eq. (8).

## III. RESULTS

Raw trajectories of the probes during a single period of oscillation are shown in Figs. 3 a-d, where for clarity, the probe displacements are magnified by a factor of 4, while their mean positions are to scale [27]. Each of the trajectories is an elliptical Lissajous figure whose major and minor axes vary systematically with angular position *θ* and distance *R* from the driven bead. It is also clear that the probe trajectories have a degree of stochasticity superimposed on their background motion.

**FIG. 3.**
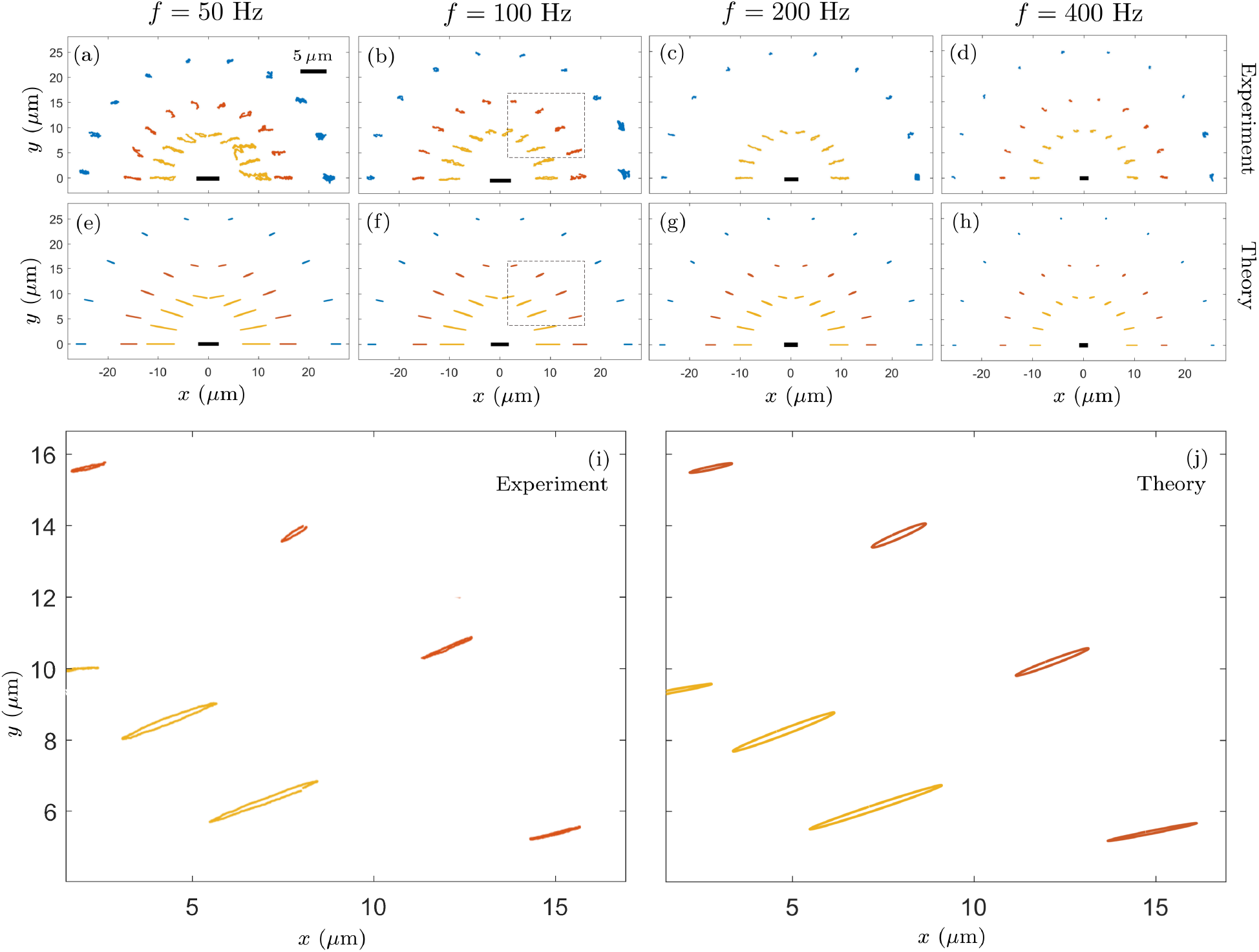
Oscillatory tracer dynamics with tracer displacements magnified by a factor 4, while their equilibrium locations are to scale. (a-d) Tracer trajectories over one period. Colors (yellow, red,blue) indicate different radial distances, as in Fig. 1. The peak-to-peak displacement of the central silica bead is indicated by the heavy black line. (e-h) Theoretical trajectories using approach given in text. (i-j) Enlarged view of trajectories in the boxed region shown in (b,h). (i) shows average cycles from experiments while (j) is simply a zoomed representation of (f). The flat elliptical trajectories, which arise from the phase shift between the components 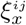 and 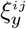 of the displacement, are accurately captured by the theory.

A quantitative treatment of the probe trajectories and an assessment of the importance of Brownian motion begin with the unsteady velocity field ***u***(***r***, *t*) due to a sphere oscillating with velocity 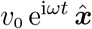. That velocity field satisfies the full Navier-Stokes equations which, if made dimensionless by the time 1/*ω*, a length *L* and velocity *U* has two Reynolds numbers [25],

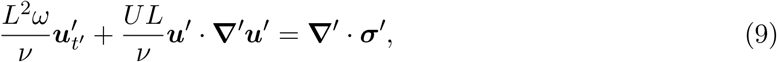

where ***σ^′^*** is the non-dimensional hydrodynamic stress. If L is on the scale of the sphere radius *a*_0_, then *U* ~ *a*_0_*ω* and *L^2^ω*/*v* and *UL/v* are both 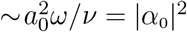, and both the inertial and nonlinear terms must be considered, but at larger length scales the time derivative dominates the nonlinear term. At the distance *R*_1_ of the closest tracers, the nonlinear term is already only ~ 10% of the inertial term and can be ignored. We have confirmed this by noting the absence of components at frequencies of 2*ω* in the power spectrum of the tracer displacements. To find the motion of the tracers we thus examine the solution of the unsteady Stokes equation *ρ_f_* ***u***_*t*_ = **∇** · ***σ***, together with the continuity equation and boundary conditions (a) 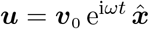 at *r* = *a*_0_ (particle surface), and ***u***_0_ → 0 as *r* → ∞. The solution, first derived by Stokes [1], can be written as ***u*** = ***u***_0_e^i*ωt*^, where

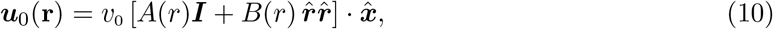

where 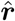 and 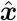 are the unit vector along the radial direction and *x*-axis respectively, and

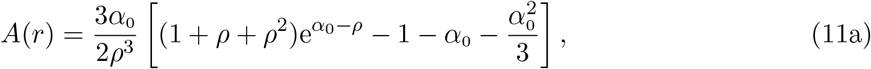

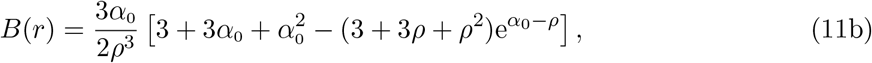

where *ρ* = (1 + *i*)*r/δ*. For comparison if the sphere were moving along the *x*-axis at a constant speed *v*_0_ the *ω* → 0 limit of (10) has the coefficients appropriate to steady flow,

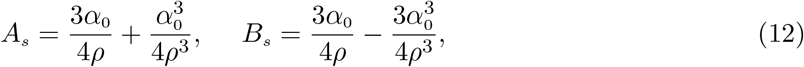

where these expressions are purely real since *α*_0_/*ρ* = *a*_0_/*r*.

We now seek the motion of a probe whose equilibrium position in the oscillating flow is **r** = (*R, θ*). As we are not referring to any particular tracer, we drop the superscript *ij*. The approach adopted here is a composite one, in which the Lagrangian inertial response of the tracer is computed from the Eulerian flow generated by the central bead. This is valid provided velocity gradients at the probe scale are small, as are the oscillation amplitude relative to the bead radius. Momentum balance for a tracer with velocity ***v***_1_e^i*ωt*^ in a fluid with velocity ***u***_0_e^i*ωt*^ takes the form [1, 26]

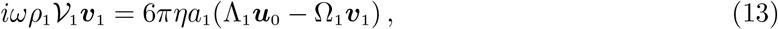

where 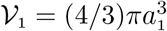 is the volume of the particle and

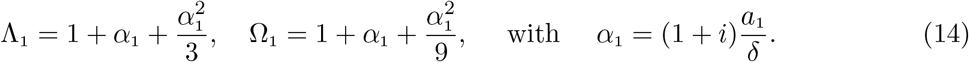

In discussing the motion of the driven particle (c.f. Eq. 3), we noted that the finite-frequency corrections to the drag law were very small; they are even smaller for tracers, whose radii are smaller by a factor of five. It follows that we may safely take Λ_1_ = Ω_1_ = 1, and thus *iωτ*_*d*1_ ***v***_i_ ≃ (***u***_0_ – **v**_1_, where in parallel with (4) we define 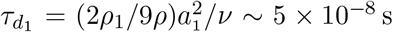. This relaxation time is so short relative to the period of driven particle oscillations that we may assume the tracer particle velocity relaxes to that of the fluid instantaneously, and thus the tracer particle velocity is simply

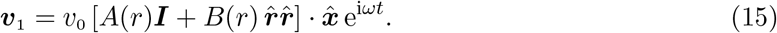

Combining this result with the response of the driven microsphere, the equations of motion for the tracer displacements, 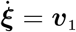, integrate to give the tracer motion at (*R, θ*),

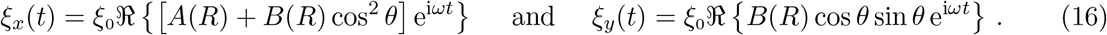

While in direct comparison with experiment we utilize the full expressions in (16), it is heuristically useful to simplify these results in the regime of low frequencies, when the distances *R_i_* of the tracers from the drive sphere are small compared to the viscous penetration depth *δ*. Thus expanding (11a) and (11b) for *α*_0_, *ρ* ≪ 1 we find

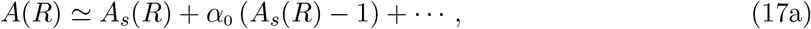

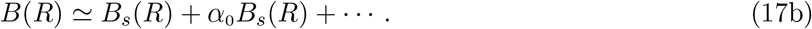

Substituting into (16), assuming as above *R/δ* ≪ 1, we observe that *A_s_*(*R*) and *B_s_*(*R*) are dominated by their Stokeslet contributions 3*a*_0_/4*R*, and thus

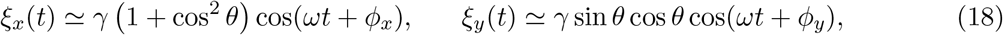

where *γ*(*R*) = (3_a0_/4*R*)*ξ*_0_, and assuming the phase shifts are small, we find

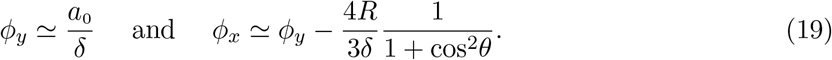

Interestingly, while *ϕ_x_* varies with the polar angle, *ϕ_y_* does not.

The Lissajous figures associated with (18) are conic sections [28], and can be rewritten as

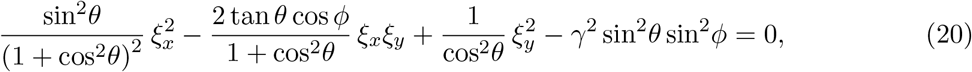

where the phase shift difference *ϕ* = *ϕ_y_* – *ϕ_x_* is

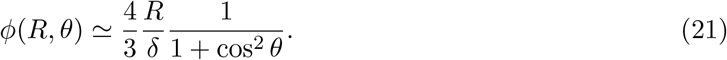

Equation 20 is in the standard form 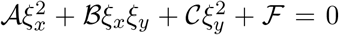 of conic sections, which are in this cases ellipses since 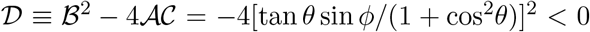. A standard analysis shows that the major axis of the ellipse is tilted with respect to the *x*-axis by an angle *ψ* satisfying 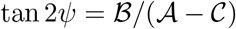. As *ψ* varies with cosϕ, corrections to the *ϕ* = 0 limit are 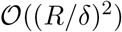, so *ψ* is well-approximated by the tilt angle of a steady stokeslet,

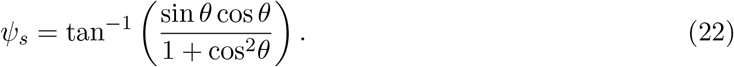

The fundamental signature of unsteadiness in the present experiment is the elliptical form of the tracer orbits. From the general expression for the semimajor and semiminor axes of ellipses,

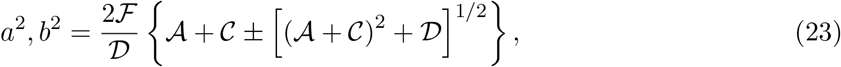

where the + (−) sign refers to *a* (*b*), we obtain the remarkably simple asymptotic results,

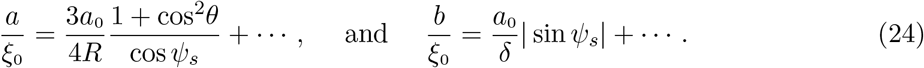

Equations 24 are the heuristic results we sought. They show that to leading order at low frequencies the semimajor axis *a* is given simply by the motion of the driven particle, projected to its position via the stokeslet contribution, while the semiminor axis *b* is nonzero only to the extent that the viscous penetration length itself is not infinite. The aspect ratios of the ellipses simply reflect the phase shift; *b/a* = (*ϕ*/2)| sin2*ψ*_s_|, and in the steady limit the ellipses degenerate into lines. For later reference we note if we include the leading unsteady corrections, then the normalized *x*-component of the tracer displacements can be written as

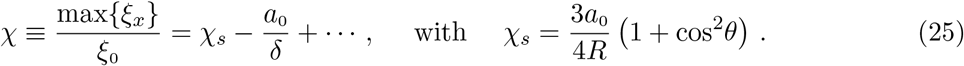

Before analyzing the tracer trajectories in detail, we use the particle trajectories to quantify the competition between the deterministic forcing of the tracer particles and thermally-driven Brownian motion. As in previous discussions of this issue [15], a useful metric with which to assess these effects is the ratio of the maximum deterministic displacement due to the oscillating fluid (the major axis of the ellipse) to the average Brownian displacement over half an oscillation period. This is essentially the square root of a Péclet number,

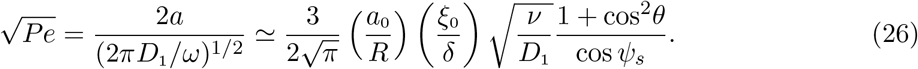

Here, *D*_1_ = *k_B_T*/6*πηa*_1_ ~ 0.4 *μ*m^2^/s (Table 1) is the diffusion constant of the tracers, with *k_B_* the Boltzmann constant and T = 298 K the absolute temperature. The last relation in (26) is obtained using the asymptotic results above and displayed to emphasize it is the product of three dimensionless ratios. The factor *v/D*_1_ is a Schmidt number Sc for the tracer particles and is very large 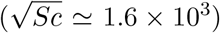, but its contribution to *Pe* is attenuated by the two small factors *a*_0_/*R* and *ξ*_0_/*δ*, each on the order of 0.1. The frequency dependence of 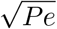 is relatively weak by virtue of the counteracting trends of *ξ*_0_ ~ *ω*^−1^ and *δ* ~ *ω*^−1/2^. Contour plots of (26) in the first quadrant of physical space where the tracers reside are shown in Figure 4, in which the semimajor axis has been computed with the full unsteady velocity field given in (10), (11a) and (11b). From these results we see that advective contributions dominate diffusion (*Pe* > 1) at all frequencies for the innermost spheres, while the two become comparable for the outermost spheres, consistent with the qualitative appearance of the trajectories.

**TABLE I.**
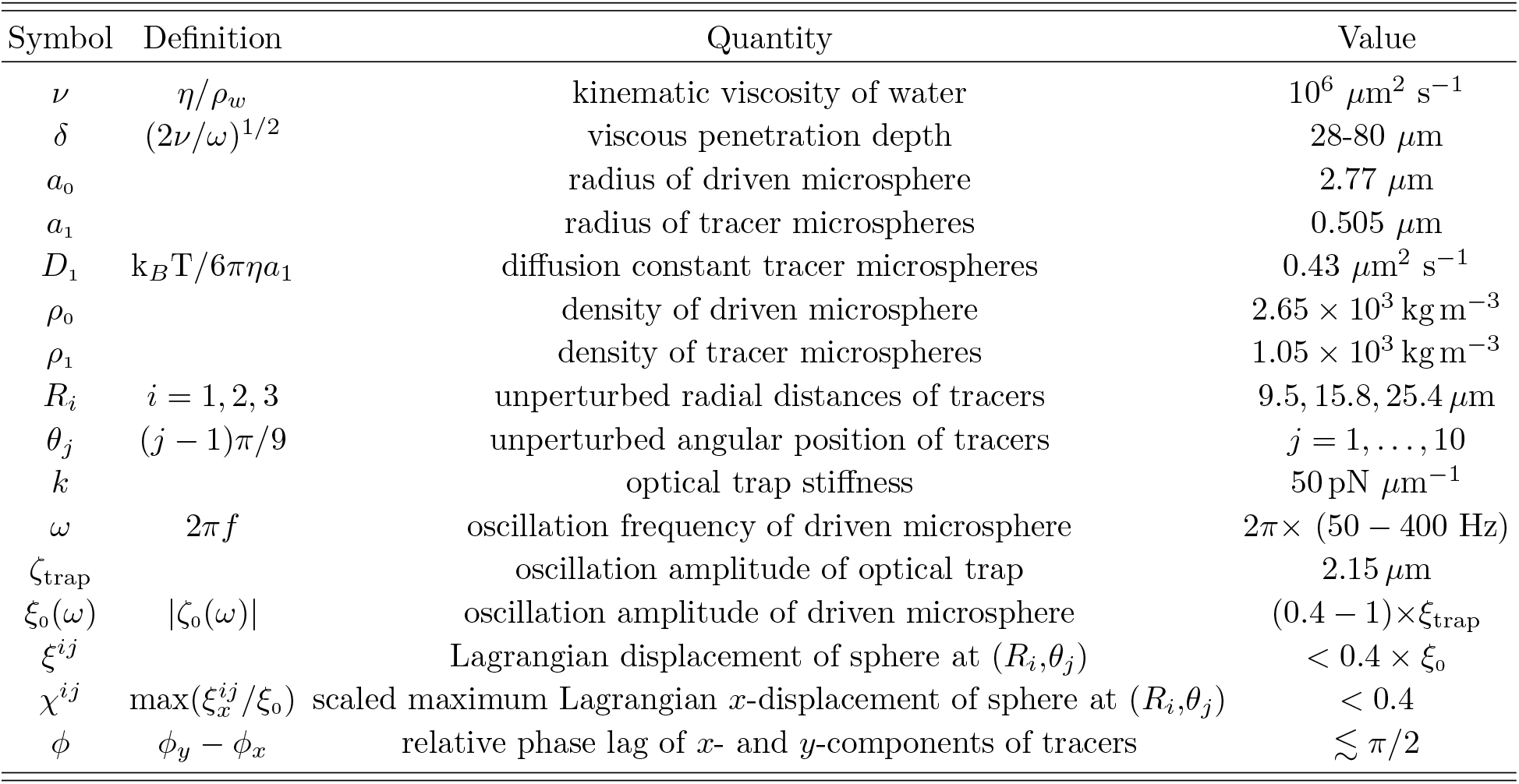
Physical quantities for experiments in water, in a convenient system of units.

**FIG. 4.**
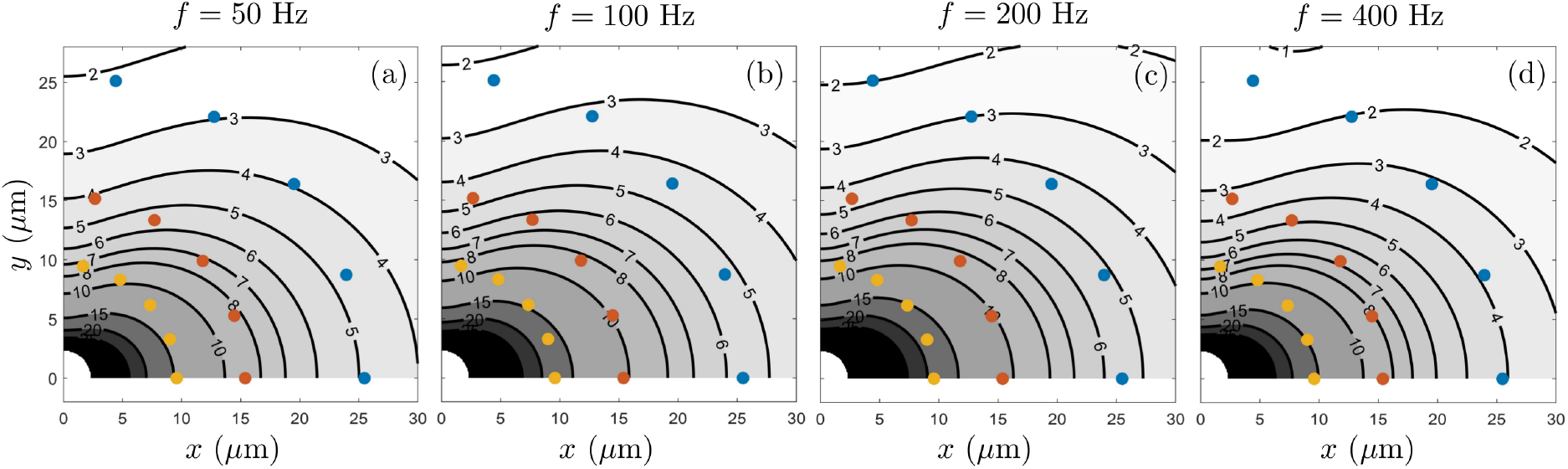
Significance of thermal fluctuations. (a)-(d) Contour plots of the 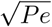 (26) in physical space, with the locations of the probe microspheres superimposed, at the four driving frequencies.

A first test of the theoretical analysis of the trajectories involves plotting the ellipses from Eq. 20 in the *x, y* plane. These are shown in Figs. 3(e-h), magnified by a factor 4 to be consistent with all the plots of the figure. In addition, the boxed portion of Fig. 3(f) is expanded in (j). In comparison, Fig. 3(i) displays the average oscillation of a few tracers at the same locations, from experiments, showing a good match with the theory. To suppress the effect of fluctuations on single oscillations, all the cycles were averaged into a composite, cyclic *x, y* path for each tracer. Each is obtained by computing the average *x*- and *y*-oscillations as a histogram with 2*πf*_*s*_/*ω* bins where *f_s_* is the sampling frequency; these data are accumulated in the bins using the time *t* mod (2*π/ω*). Alternatively, Fig. 5 shows two basic geometrical features of the elliptical tracer trajectories, their orientation and major axis, each expected to be dominated by their steady contributions. The orientation angle in Fig. 5(a) agrees well with the steady angle *ψ_s_* in (24), and the semi-major axis (Fig. 5(a)) is likewise well described by the leading order relation in (22).

**FIG. 5.**
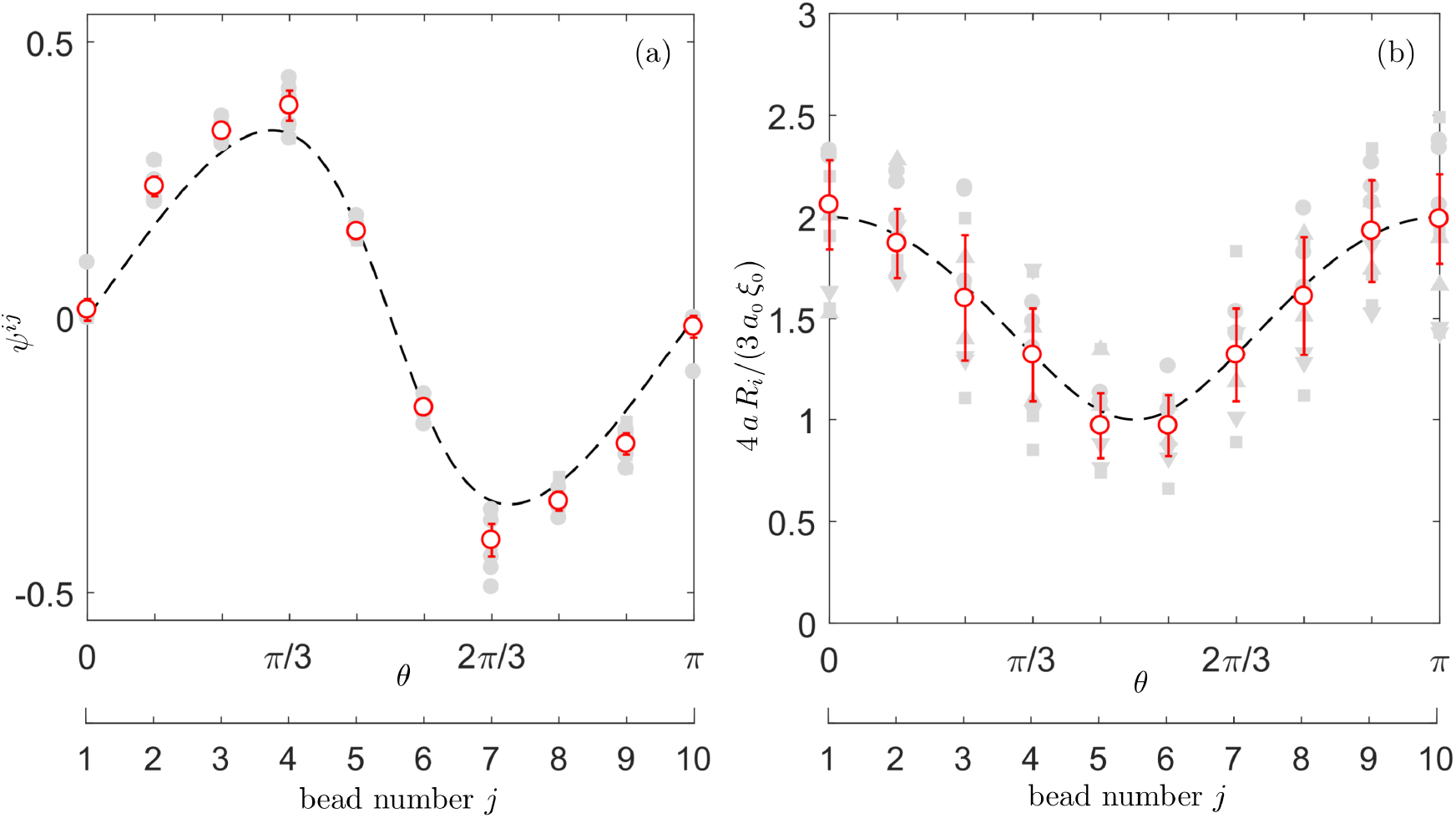
Orientation and size of elliptical tracer trajectories. (a) Tilt angle of the major axis of elliptical Lissajous figures as a function of mean angular position relative to driving axis. Gray symbols represent the individual data points at the four experimental frequencies (*f* = 50, 100, 200, and 400 Hz) and three radii (*R* = 9.5, 15.8, and 25.4 *μ*m) at a given angular position *θ_i_*. Open red circles denote the mean values of each of those measurements at a given θ and their standard deviations. Dashed line is the steady tilt angle (22). (b) As in (a), but for the semimajor axis of the ellipses. Dashed line is the low-frequency limit (24).

Focusing on the displacements along the same (*x*) axis as the driven microsphere, Fig. 6 summarizes the results for the amplitudes and phase shifts of the tracers. In (a-d) we plot the normalized component of the displacement as defined in (25),

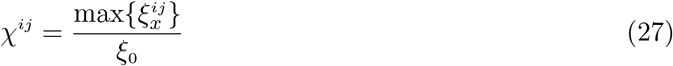

for experiments (symbols) and theory (lines). In the experiments, the relative amplitude and phase compared to the driven bead are measured from the fast Fourier transform (FFT) of the *x*(*t*) data of each tracer, by identifying the amplitude (respectively, the phase) from the peaks in the magnitude (phase) plots of the FFT for the tracers and dividing by the magnitude (or subtracting the phase) from the FFT of the driven bead. At any given frequency the agreement between the data and the steady theory (shown by dashed lines) is best for those tracers closest to the driven particle and progressively decreases for more distant probes, while at any given radius the agreement with the steady theory worsens at higher frequencies. For example, the deterministic component of the displacement is overestimated by about 20% for *f* = 50 Hz, and up to 100% (for *f* = 400 Hz) for the most remote ones - i.e at a distance *R*_3_ from the origin. Both of these trends are fully consistent with the relevant measure of unsteadiness being *R/δ*. In Fig. 6b we show the the experimental phase shift *ϕ_j_* between the tracers and the active particle. The magnitude and angular dependence are both accurately captured by the unsteady theory. At the very highest frequency used, the phase shifts of the most distant probes located close to the *y*-axis —at (*R*_3_, *θ*_5_) and (*R*_3_,*θ*_6_)—are very large; the probes are almost in quadrature with the forcing. We see from these results that despite a large displacement of the central bead (which is of the same order as its radius) and a direct use of the Eulerian form of the viscous unsteady flow ***u***_0_, the agreement between theory and experiment is very good, with a maximum relative error of 4% overall.

**FIG. 6.**
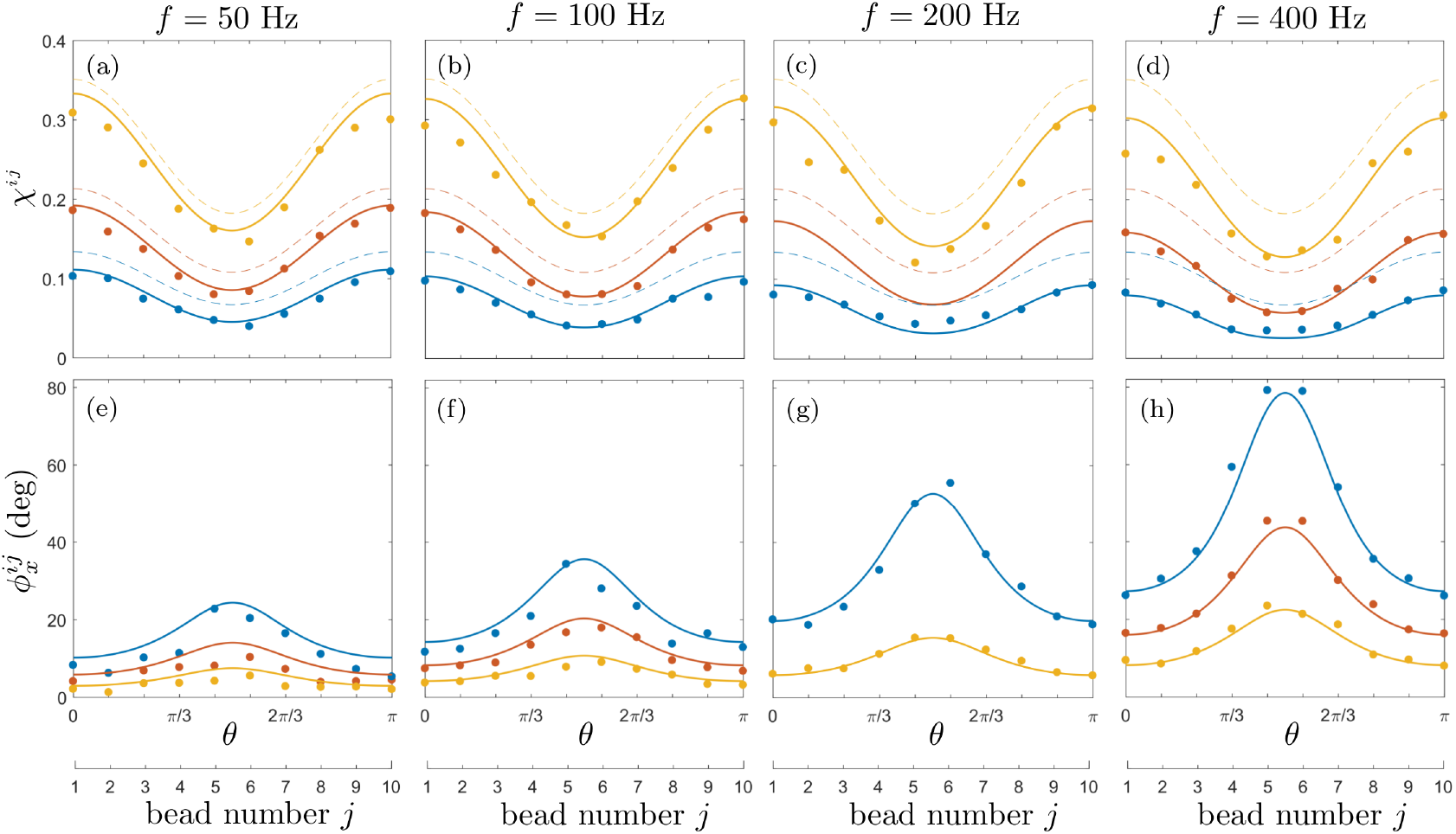
Dynamics of the passive tracers. (a-d) Normalized maximum displacements (solid circles), and their theoretical counterpart *χ* (solid lines). Dashed line show displacements calculated within steady Stokes equation. (e-h) Phase shifts between responses of the probes and the harmonic motion of driving bead (solid circles), and theoretical counterpart (solid lines).

As a final test of the unsteady theory, we ask whether the data in Fig. 6 are consistent with the predicted leading-order low-frequency limits in the sense of a data collapse. Focusing on the same *x*-component of the phase shift and amplitude, the analysis in (19) and (21) can be written the scaling forms for the phase and amplitude,

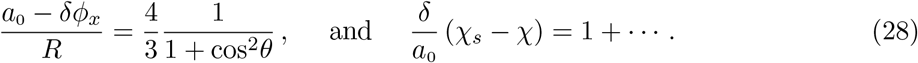

Figure 7 shows good agreement in both cases, especially for the phase shift.

**FIG. 7.**
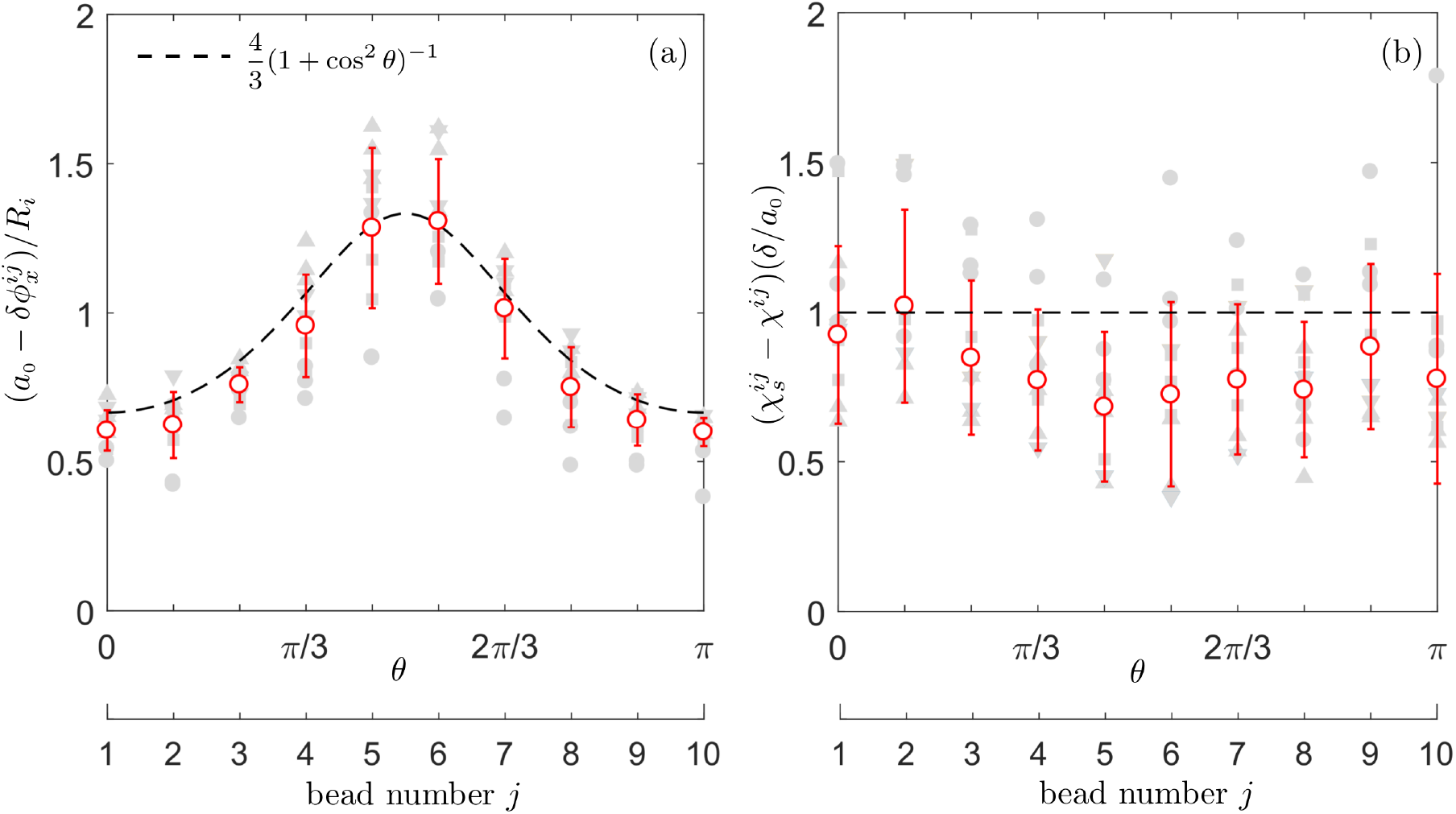
Test of the low-frequency scaling. (a) Rescaled phase shift as a function of angle, using same symbol motif as in Fig. 5. Dashed line is the scaling result (28). (b) As in (a), but for the oscillation amplitude, with dashed line from (28) to leading order. There are no adjustable parameters in (a) or (b).

## IV. DISCUSSION

We have shown here how optical trapping and particle-tracking techniques allow for a precise microscale test of the theory of unsteady Stokes flows. At the scale of colloidal particles, and with oscillation frequencies in the range found in biological systems, the simplifications arising from lack of inertial effects on particle motion and corrections to the Stokes drag law allow for a simple and compact picture of the particle orbits. The regime of sizes and frequencies explored is also such that Brownian motion makes only a modest contribution to the tracer dynamics, with an effective Péclet number generally exceeding unity. Our experimental results show that tracer particles move on simple elliptical orbits even in a regime with very large phase shifts, in quantitative accord with a low-frequency analysis. These experimental observations would be difficult to reproduce by conventional particle imaging techniques which are based on obtaining an Eulerian velocity map from correlation functions of small tracer displacements.

As outlined in the introduction, one clear motivation for the present study is provided by the evidence that unsteady effects are present during the collective beating of eukaryotic cilia and flagella. It is an open question as to whether these effects actually control synchronization. Two features of the present work will likely bear on this issue; the angular dependence of the phase shift and the elliptical orbits themselves. The former dictates the strength of the lateral coupling between cilia along a tissue, while the latter represents vorticity created by the driven particle that may be relevant to wave propagation. It is important to note that in the many cases in which metachronal waves occur there is a nearby underlying no-slip surface —the cell wall of a ciliate, or the tissue surface of an ciliated epithelium —whose presence can not be ignored. Indeed, a recent study of synchronization in arrays of “colloidal oscillators” [14], microspheres moved along periodic orbits by optical traps, show that surface proximity can profoundly affect the collective dynamics that they exhibit. Thus, a natural next step is the study of model unsteady flows near no-slip surfaces [29].

## V. ACKNOWLEDGEMENTS

This work was supported in part by ERC Consolidator grant 682754 (EL), ERC PoC grant CellsBox (PC and JK), Wellcome Trust Investigator Award 207510/Z/17/Z, Established Career Fellowship EP/M017982/1 from the Engineering and Physical Sciences Research Council, and the Marine Microbiology Initiative of the Gordon and Betty Moore Foundation, Grant 7523 (REG).

